# Time-tagged ticker tapes for intracellular recordings

**DOI:** 10.1101/2021.10.13.463862

**Authors:** Dingchang Lin, Xiuyuan (Ted) Li, Pojeong Park, Benjamin Tang, Hao Shen, Jonathan B. Grimm, Natalie Falco, David Baker, Luke D. Lavis, Adam E. Cohen

**Affiliations:** Department of Chemistry and Chemical Biology, Harvard University, Cambridge, MA 02138; University of Washington, Molecular Engineering and Sciences, Seattle, WA 98195; Janelia Research Campus, Howard Hughes Medical Institute, 19700 Helix Drive, Ashburn, VA 20147; Department of Physics, Harvard University, Cambridge, MA 02138

**Author notes:** equal contribution (Dingchang Lin, Xiuyuan (Ted) Li).

## Abstract

A core taken in a tree today can reveal climate events from centuries past. Here we adapt this idea to record histories of neural activation. We engineered slowly growing intracellular protein fibers which can incorporate diverse fluorescent marks during growth to store linear ticker tape-like histories. An embedded HaloTag reporter incorporated user-supplied HaloTag-ligand dyes, leading to colored stripes whose boundaries mapped fiber growth to wall-clock time. A co-expressed eGFP tag driven by the *cFos* immediate early gene promoter recorded the history of neural activity. High-resolution multispectral imaging on fixed samples read the cellular histories. We demonstrated recordings of *cFos* activation in ensembles of cultured neurons with a single-cell absolute accuracy of approximately 39 min over a 12-hour interval. Protein-based ticker tapes have the potential to achieve massively parallel single-cell recordings of multiple physiological modalities.

## Introduction

Optical imaging typically faces a tradeoff between temporal and spatial information: one can image vast numbers of cells in fixed tissue,^1^ but to record dynamical information requires live-tissue optical access, in which the field of view is limited by light scatter and optical instrumentation. These challenges are particularly evident when trying to image markers of brain activity, such as activity-responsive immediate early genes (IEGs), where relevant dynamics are broadly distributed throughout the brain.^2^ Tools to record the dynamics of large numbers of cells, without constraints from *in vivo* imaging, could transform our ability to study ensemble dynamics in the nervous system and in other tissues.

Reporter genes or antibody labels can report brain-wide patterns of IEG activation, but typically at only one^3,4^ or at most two^5^ time-points. Time-lapse *in vivo* microscopy can report longitudinal IEG dynamics,^6^ but only in a small optically accessible region. Theoretical analyses have explored the possibility of encoding brain-wide dynamics in DNA or RNA sequences,^7,8^ but despite progress^9^ such ideas have not yet been realized.

Natural histories are written in the patterns of tooth enamel, the structures of pearls, and the thickness of tree rings. Inspired by these natural phenomena, we sought to encode cellular histories in protein microcrystals within individual cells. Protein assemblies can last for months or years and offer a wide array of functionalities which could serve as the basis for recording schemes.

A protein-based recording scheme must have three elements (Fig. 1). First, it requires a protein scaffold which grows with time and which can incorporate fluorescent marks. Second, it requires a means to impart fiducial timestamps to relate scaffold growth (which will likely vary between cells and over time) to timing of events in the outside world. Third, it requires a fluorescent reporter of cellular activity which can be stably incorporated into the scaffold.

**Figure 1.**
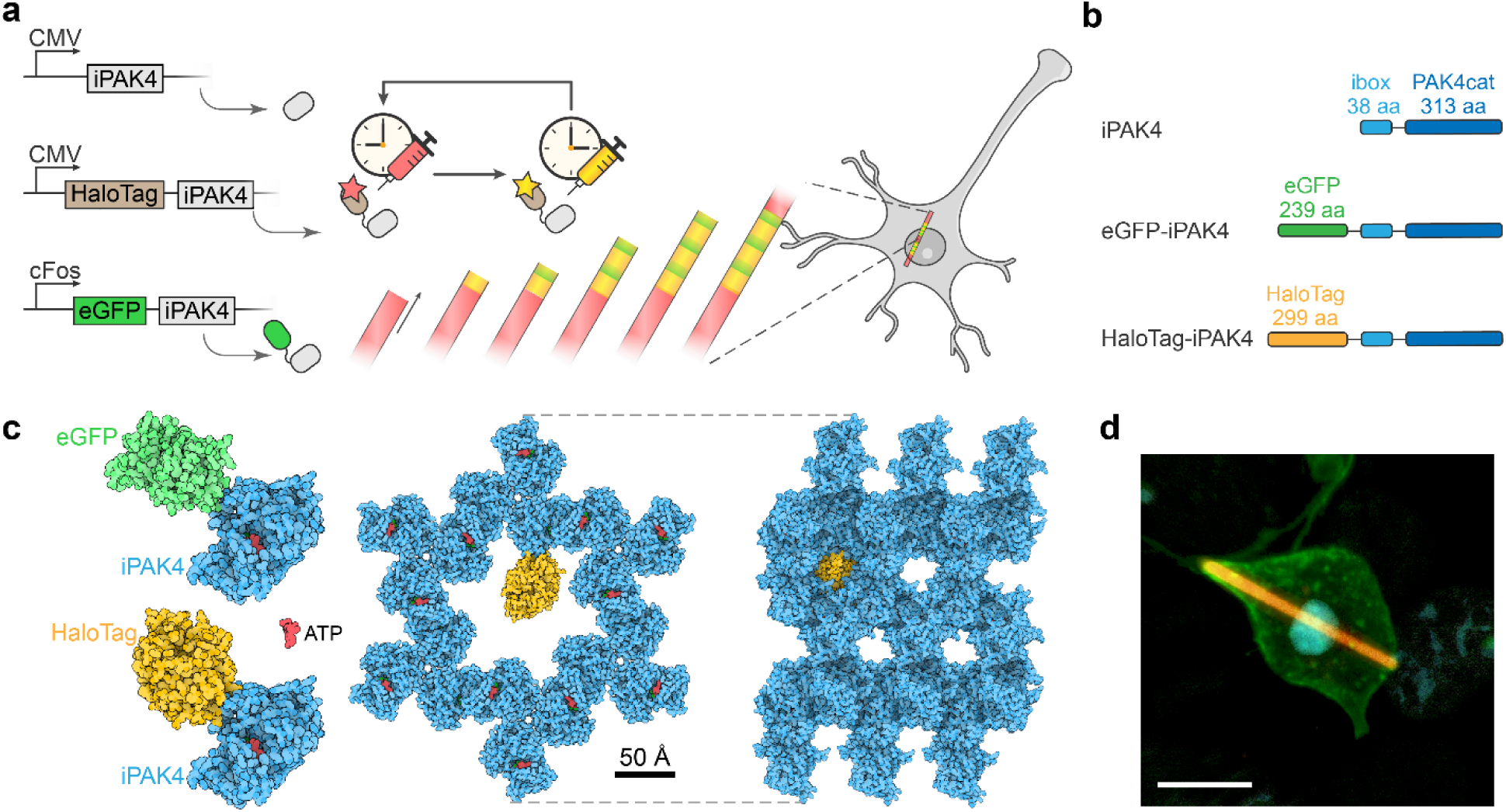
iPAK4 forms intracellular protein fibers. A) Scheme for intracellular recording of cFos activity with fiducial timestamps. iPAK4 forms the fiber scaffold. HaloTag-iPAK4 incorporates HT dyes, permitting labeling of the fiber with fiducial timestamps. Neural activation drives expression of cFos::eGFP-iPAK4, introducing green bands into the fiber. B) Composition of the protein constructs used to label intracellular protein fibers. C) The structures of tagged iPAK4 monomers and the crystal structure with hexagonal pores (from PDB:4XBR) were modeled using Protein Imager software._38_ D) Image of a HEK cell expressing CMV::iPAK4 (95%) and CMV::HT-iPAK4 (5%). The fiber was stained with JFX_608_, the membrane was labeled by expressing GPI-eGFP, and the nucleus was labeled with DAPI. Scale bar 10 μm.

For the protein scaffold, our selection criteria were that it should: express in mammalian cells, assemble into a growing structure from a single polypeptide, have a known crystal structure, be unlikely to interfere with cellular physiology, and accommodate decoration with a fluorescent tag without disrupting the structure of the assembly. We considered many possibilities, including endogenous microtubules, bacterial R-bodies,^10^ plant forisomes,^11^ amyloid fibrils,^12^ prions,^13^ filamentous viruses,^14^ crystals of the fluorescent protein XpA,^15^ engineered fiber-forming peptides,^16^ and other proteins that spontaneously crystallize in cells.^17^

A fusion of the catalytic domain of the Pak4 kinase and its inhibitor Inka1 (hence called iPAK4) largely satisfied the selection criteria. This construct has been shown to stably assemble in cells into rod-shaped crystals.^18^ Remarkably, the crystal structure has a hexagonal array of internal pores large enough to accommodate eGFP or a HaloTag, suggesting the possibility of linear encoding of information via patterned fluorescence (Fig. 1c).

We sought to use a fusion of the HaloTag (HT) to iPAK4 to provide fiducial timestamps. We reasoned that washes with different colored HT-ligand dyes could create color boundaries whose positions would correspond to known times. Even though HT dye washout *in vivo* occurs over hours, HT dye injection and labeling have fast onset (< 10 minutes *in vivo*),^19^ permitting precise demarcation of timestamps by the location of a color transition. A broad palette of bright, photostable, and brain-penetrant Janelia Fluor HT-ligand dyes are available, permitting diverse spectral encodings of fiducial timestamps. ^20–22^

The activity reporter should store a stable mark of cellular activity or physiology upon incorporation into the growing fiber. IEG reporters are a powerful tool for identifying the brain regions and neuronal subtypes activated in a particular context.^23^ For example, the IEG *cFos* has been used to map neurons activated during feeding,^24^ sleep,^25^ parenting,^26^ or aggression.^27^ IEG activity is also used to identify the neurons recruited into the physical embodiment of a memory trace, or engram. Motivated by potential applications toward brain-wide memory mapping, we used the *cFos* promoter to drive expression of eGFP-iPAK4.

## Results

In HEK cells co-transfected with CMV::iPAK4 (95%) and CMV::HT-iPAK4 (5%), fibers began to grow 14-20 h after transfection. Incubation with a HT-ligand dye made the fibers brightly fluorescent (Fig. 1D, Methods). Most fiber-containing cells had only one fiber (288 of 333 cells, 86%). When the fibers grew longer than the cell diameter, the membrane deformed around the fiber, though this did not appear to impair either fiber growth or cell viability (Fig. S1a).

To assess the suitability of iPAK4 fibers as a recording medium, we performed a detailed characterization of their nucleation and growth. We co-expressed CMV::iPAK4 (95%) and CMV::eGFP-iPAK4 (5%) in HEK cells and recorded time-lapse video microscopy over a 13 h interval starting 10 h after transfection (Methods, Movie S1). We then tracked the fluorescence of the cytoplasm and the growth of individual fibers (Methods, Movie S2). Initially, green fluorescence accumulated in the cytoplasm. Upon nucleation, fibers grew quickly (∼0.5 μm/min) at first, while the cytoplasmic fluorescence dropped. Fiber growth then slowed to a constant rate of 1-2 μm/h, suggesting a balance of protein translation and fiber growth (Fig. S1b). We ascribe the rarity of multiple fibers per cell to the fact that a single growing fiber maintained the soluble iPAK4 concentration below the nucleation threshold.

To test whether fibers could encode HT dye timestamps, we successively washed HEK cells growing HT-iPAK4 fibers with different colors of HT-ligand dyes at Δt = 2 h intervals in the sequence JF_503_, JF_669_, JFX_608_, JF_503_, JF_669_ (Fig. 2a; see Fig. S2 for dye structures and photophysical properties). We then fixed the cells and imaged with spectrally resolved confocal microscopy (Methods). We observed clear progression of colored bands matching the sequence of dye additions. The bands followed mirror-image patterns on opposite sides of the fibers, indicating that the fibers grew from both ends (Fig. 2b, S3).

**Figure 2.**
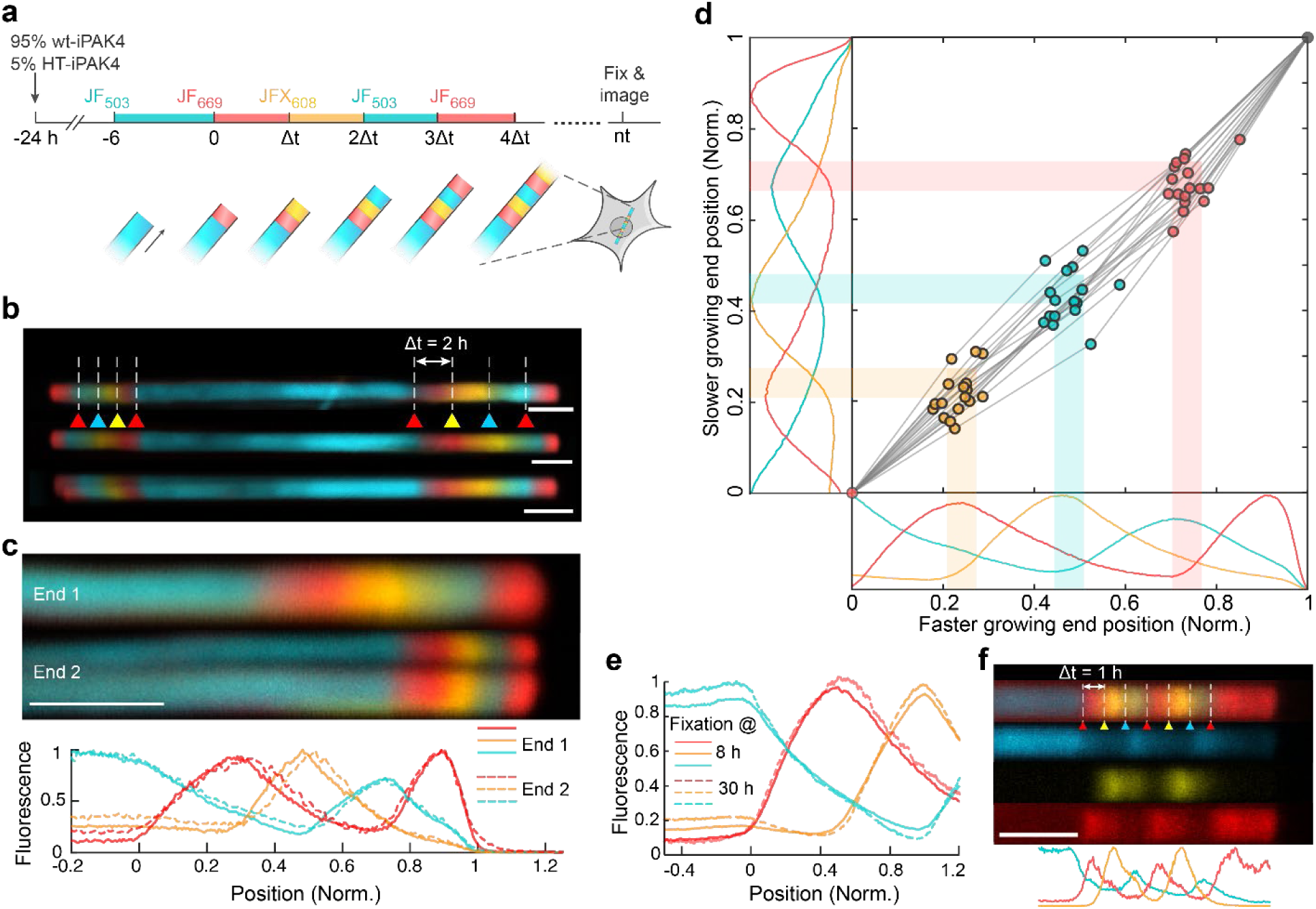
iPAK4 fibers can be sequentially labeled with different dyes. A) Scheme for multi-color labeling of intracellular iPAK4 fibers. B) Images of iPAK4 fibers labeled with four dye transitions on each end. Triangles indicate timing of dye addition. Blue: JF_503_, Red: JF_669_, Yellow: JFX_608_. Scale bars 5 μm. C) Comparison of two ends of a single fiber. Top: images of the two ends. In this case, the fiber split at one of its ends, an occurrence in approximately 10% of fibers. Bottom: quantification of the fluorescence traces in the two ends, normalized to the first JF_669_ addition and the end of the fiber. Scale bar 5 μm. D) Comparison of dye transition points on the faster and slower-growing fiber ends. The scatter plot shows the positions of the JFX_608_, JF_503_ and JF_669_ transitions normalized relative to the first JF_669_ addition (at position 0) and the end of the fiber (at position 1). The plots show the mean fluorescence profiles of *N* = 22 fibers. E) Mean fluorescence profiles of fibers exposed to the same sequence of dyes and then grown for different amounts of time (*N* = 22 fixed at 8 h, 12 fixed at 30 h). The overlap of these profiles indicates negligible monomer exchange over 22 h. F) Fiber profile with seven dye switches separated by 1 h. Scale bar 5 μm.

Dye additions appeared as upward-going kinks in the fluorescence profiles (Fig. S3), which we located as peaks in the second derivative of fluorescence vs. position (Methods). By dividing the transition zone widths (full-width at half-maximum of the peak in the second derivative) by the fiber-specific growth rates, we calculated a mean transition zone duration of 7.5 min (Fig. S4a).

To determine the influence of HT dye labeling kinetics on the widths of these transitions, we measured the labeling of intracellular HT receptors with different HT dyes (Methods). These measurements probed the combined effects of membrane permeation and the HT reaction. The effective time constants at 1 μm dye were: 43 min (JF_503_), 3.5 min (JF_525_), 2.0 min (JF_552_), 12 min (JFX_608_), and 17 min (JF_669_) (Fig. S4b). The precision of localizing a dye transition in a fiber is not equal to the labeling time constant, but rather the precision with which one can identify the onset of the labeling reaction. For all the JF dyes, this time was much less than 10 min (Fig. S4b). Together, these results established that HT dye transitions provided a means to timestamp iPAK4 fiber growth with a local precision (i.e. around the time of the time stamp) of ∼10 min or better.

Due to the P6_3_ symmetry of the iPAK4 crystal structure,^18^ the two ends are not chemically equivalent and need not have the same growth rate. However, the asymmetry in growth rate was modest, with the slower end growing, on average, at 62 ± 19% of the speed of the faster end (mean ± s.d., *N* = 17 fibers, Fig. S5). We then compared the fluorescence profiles on the two ends. To account for the difference in growth rate between the fiber ends, we mapped the first dye addition (JF_669_ at t = 0) to x = 0, and the end of the fiber (fixation at t = 8 h) to x = 1. After normalizing the spatial scales, the fast and slow-growing fiber ends showed very similar profiles (Fig. 2c).

To test whether both fiber ends could be useful for recording fiducial timestamps, we analyzed the locations of the dye transitions on both ends of *N* = 17 fibers. For both the faster and slower-growing ends, the normalized locations of dye transitions at *t* = 2, 4, and 6 h mapped linearly onto position between the first dye addition (t = 0) and the end of the fiber (t = 8 h) (Fig. 2d). The standard deviations in the inferred timing (averaging over the three transitions) were 18.3 min on the fast end and 24.8 min on the slow end. Together, these results established that both ends of the fibers could record fiducial timestamps with an absolute accuracy of better than 25 min over an 8 h baseline.

If the residual errors in timing were driven by a factor shared by the two fiber ends (e.g. variations in iPAK4 expression level), then these errors would lie primarily along the diagonal line *Position 1* = *Position 2*. We calculated the cross-correlation in the timing errors between the two fiber ends, and averaged over all transitions (*t* = 2, 4, and 6 h) and all fibers. This cross-correlation was only 0.32, implying that the variations in growth rate were primarily driven by factors local to each end. In principle, timing precision could be improved by averaging measurements on the two fiber ends, but we found that often the image quality was better on one end than the other because of differences in focal plane or presence of out-of-focus fibers. Consequently, we typically analyzed only one fiber end per cell, and we did not attempt to distinguish between the faster- and slower-growing ends.

A protein ticker tape should stably store its information for extended times. Since the iPAK4 fibers were held together by non-covalent interactions, we tested whether monomer exchange blurred the HT dye boundaries over time. Two dishes were exposed to the same sequence of dye switches at intervals of Δt = 2 h. One dish was fixed at *t* = 8 h, and the other was returned to the incubator and fixed a day later at *t* = 30 h. We then compared the fluorescence profiles for the section of the fibers that grew concurrently (from *t* = 0 to 8 h). The mean profiles were nearly indistinguishable between the early dish (*N* = 22 fibers) and the late dish (*N* = 12 fibers) (Fig. 2e). Thus, monomer exchange was negligible over one day.

To test the limits of how fast we could encode dye transitions, we made fibers with seven dye switches at Δt = 1 h. At this short interval peaks were still visible, but their amplitude was suppressed relative to the parts of the fiber with no dye switches, indicating incomplete transitions in dye labeling. Analysis of the dye profiles for more widely spaced dye transitions indicated a half-life of soluble HT-iPAK4 of 4.5 h (Fig. S4a), explaining the loss of signal at faster dye switches. These results indicated that fiducial timestamps should be separated by at least Δt = 2 h.

We next studied the precision with which the timing of cellular events could be identified. If each fiber end grew at a constant rate, the timing of cellular events could be linearly interpolated between timestamps with a precision far greater than the interval between the timestamps. However, variations in growth rate might degrade the precision. To determine the precision empirically, we incubated fibers in JF_525_, and then switched to JF_669_ at *t* = 0. In different dishes we doped in the dye JFX_608_ at *t* = 2, 4, 6, 7, 8, or 9 h to simulate the onset of a cellular event. Finally, we switched back to JF_525_ at *t* = 10 h, grew the fibers for another 10 h, and fixed the dishes at *t* = 20 h (Fig. 3a).

**Figure 3.**
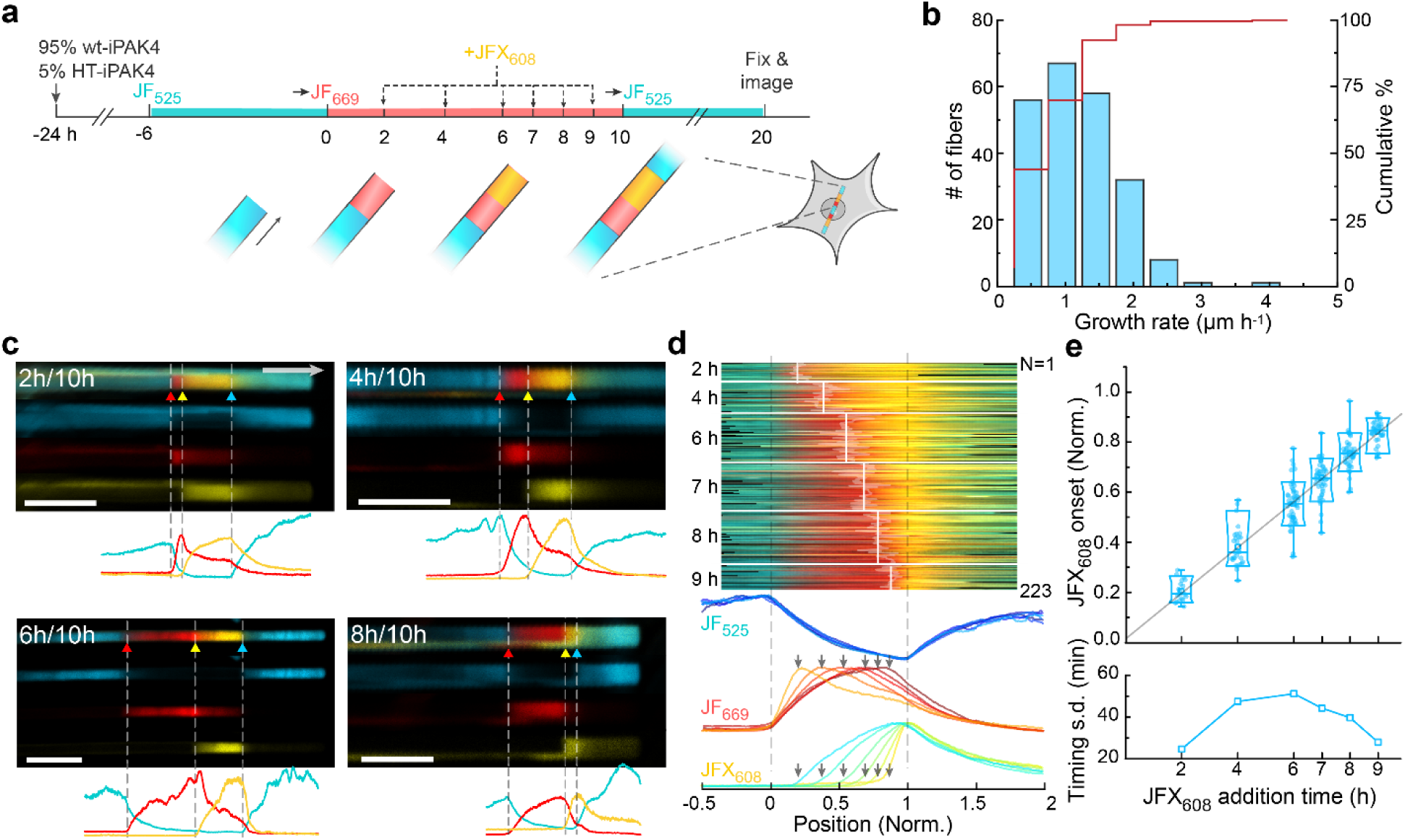
iPAK4 fibers report timing of intracellular events. A) Experimental design for testing the precision with which addition of JFX_608_ could be determined relative to timestamps from addition of JF_669_ (at *t* = 0) and JF_525_ (at *t* = 10 h). B) Histogram of growth rates between the timestamps at *t* = 0 and *t* = 10 h. C) Top: Images of fibers with JFX_608_ addition at *t* = 2, 4, 6, and 8 h. Bottom: fluorescence line profiles. Scale bars 10 μm. D) Top: fluorescence profiles of *N* = 223 fiber ends with JFX_608_ addition at different timepoints. The profile lengths have been normalized to line up the timestamps at *t* = 0 and 10 h. Bottom: Mean fluorescence traces in each of the three dye color channels, for all times of JFX_608_ addition. E) Top: positions of the JFX_608_ onset as a function of dye addition time. Bottom: standard deviation in the inferred timing of JFX_608_ addition for each population of fibers.

Over the 10 h interval between addition of JF_669_ and return to JF_525_, the fibers grew at 1.1 ± 0.58 μm/s (mean ± s.d., *N* = 223 fibers, Fig. 3b). Visual inspection of the fibers with JFX_608_ addition at different timepoints showed clear onset of yellow staining at the corresponding positions along the red band (Fig. 3c). We quantified the three-color fluorescence profiles of *N* = 223 fibers (20 to 51 fibers per time-point of JFX_608_ addition) and identified the locations of the three dye additions (JF_669_, JFX_608_, JF_525_). As above, we mapped each fluorescence trace so that the first and third dye additions occurred at x = 0 and 1, respectively (Fig. 3d).

The mean traces clearly showed the time-dependent onset of JFX_608_ fluorescence, which was also evident in low-magnification images of the fiber population (Fig. S6). We then quantified the distributions of JFX_608_ onset. The distribution linearly mapped dye addition time to normalized position between 0 and 1 (Fig. 3e). Mapping the standard deviation of JFX_608_ labeling onset onto the 10 h time axis yielded single-fiber precisions between 25 and 51 min (Fig. 3e). The precision was greatest near the fiducial points at *t* = 0 and 10 h, and lowest in the middle of the trajectory. This observation established that the uncertainty in JFX_608_ timing was dominated by nonuniformity in the fiber growth rate as opposed to errors in locating the JFX_608_ transition points, since localization errors would be statistically similar anywhere along the fiber. Thus to achieve greatest temporal precision for detecting an event, one should deposit a fiducial timestamp near the candidate event.

Finally, we tested the ability of iPAK4 ticker tapes to report activation of the IEG cFos in cultured rat hippocampal neurons. We first confirmed the ability of switches in HT dye labeling to impart fiducial time stamps on HT-iPAK4 fibers in neurons (Fig. S7). We then tested whether iPAK4 fibers affected neuronal electrophysiology. Neurons infected with lentivirus encoding CMV::iPAK4 (90%) and CMV::eGFP-iPAK4 (10%) grew single straight fibers (Fig. 4a). We used patch clamp recordings to compare the electrophysiology of neurons with and without fibers. Although in many cases the fibers length was several times longer than the soma diameter, neurons with fibers spiked normally (Fig. 4b), and had membrane resistance, membrane capacitance, resting potential, and rheobase which were statistically indistinguishable from neurons without fibers (*N* = 11 neurons with fibers, *N* = 12 without, Fig. 4c).

**Figure 4.**
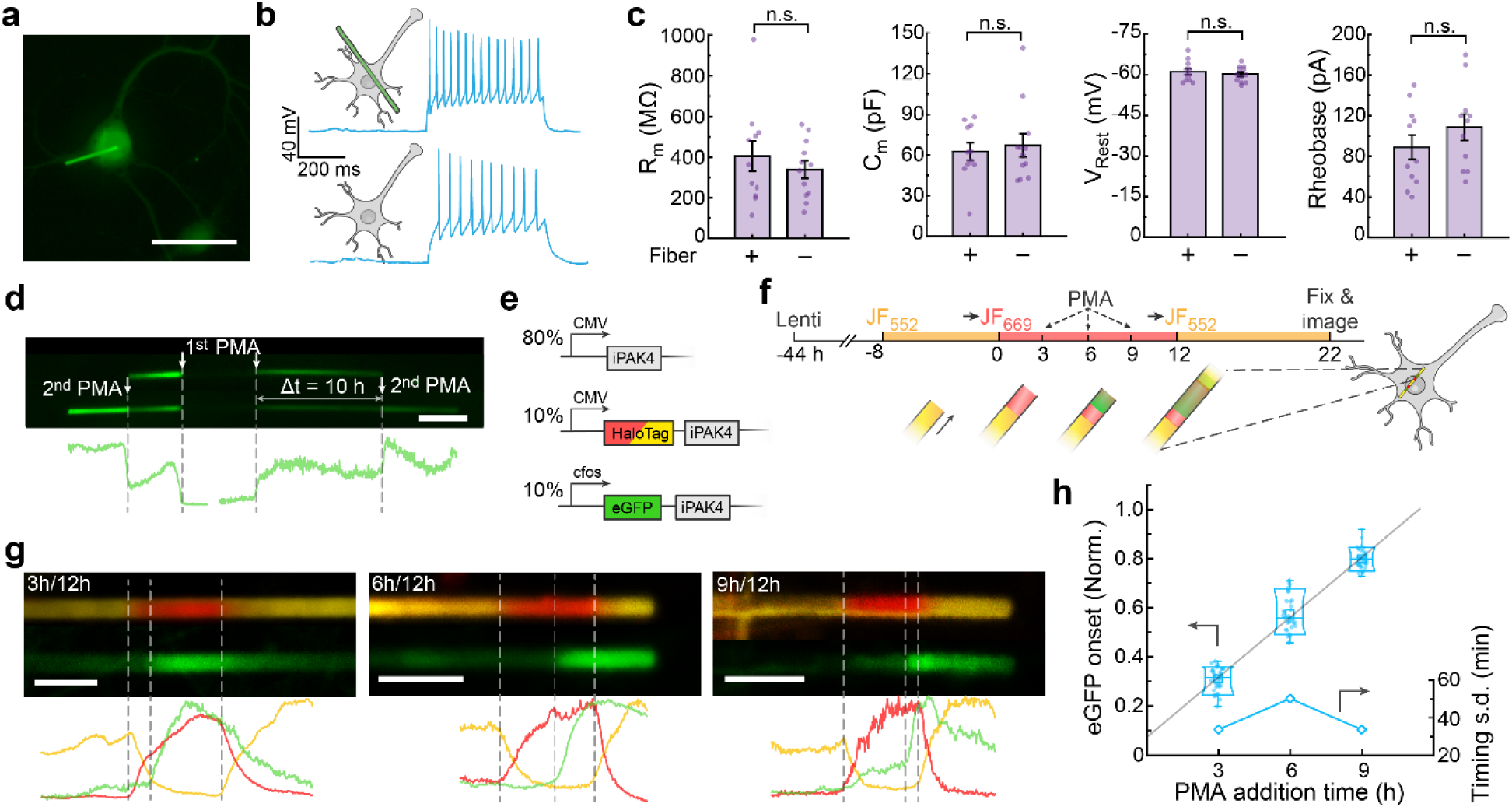
Protein ticker tape recordings of cFos activation in neurons. A) Image of a cultured neuron expressing lentiviral CMV::iPAK4 (90%) and CMV::eGFP-iPAK4 (10%). Scale bar 50 μm. B) Representative patch clamp recordings in neurons with or without an iPAK4 fiber. Spikes were evoked by a current injection of 100 pA. C) There were no significant differences between neurons with or without fibers in membrane resistance (404 ± 73 MΩ vs 339 ± 43 MΩ, *p* = 0.43), membrane capacitance (63 ± 6 pF vs 67 ± 9, *p* = 0.67), resting potential (−61.2 ± 1.2 mV vs −60.2 ± 0.8 mV, *p* = 0.48), or rheobase (89 ± 12 pA vs 109 ± 13 pA, *p* = 0.27, *N* = 11 neurons with fibers, 12 neurons without). D) Fiber in a neuron expressing lentiviral CMV::iPAK4 (90%) and cFos::eGFP-iPAK4 (10%). The top image shows the fiber after a first PMA addition, the bottom image shows the same fiber after a second PMA addition. Scale bar 5 μm. E) Genetic constructs for recording time-tagged cFos activation in neurons. F) Experimental protocol for recording time-tagged cFos activation in neurons. Transitions to JF_669_ at *t* = 0 and to JF_552_ at *t* = 12 h provided fiducial timestamps. cFos was activated via addition of PMA at *t* = 3, 6, or 9 h. G) Top: representative images of fibers with PMA addition at *t* = 3, 6, or 9 h. Bottom: fluorescence line profiles. Scale bar 5 μm. H) Normalized positions of eGFP onset relative to fiducial timestamps at *t* = 0 and 12 h.

In neurons co-expressing CMV::iPAK4 (90%) and cFos::eGFP-iPAK4 (10%), fibers initially grew with little green fluorescence. Addition of phorbol 12-myristate 13-acetate (PMA, 1 μm), an activator of cFos expression,^28^ led to bands of bright green fluorescence (Fig. S8). Sequential additions of PMA to a single dish led to distinct bands of eGFP fluorescence, clearly resolved by sharp eGFP boundaries (Fig. 4g).

To measure timing of cFos activation relative to fiducial timestamps, we co-expressed CMV::iPAK4 (80%), CMV::HT-iPAK4 (10%), and cFos::eGFP-iPAK4 (10%). We stained the neurons with JF_552_, and then switched to JF_669_ at *t* = 0. We then added PMA (1 μm) at *t* = 3, 6, or 9 h, and introduced a second fiducial timestamp by switching back to JF_552_ at *t* = 12 h. Fibers were grown until *t* = 22 h and then fixed and imaged.

We observed clear green bands indicating cFos-driven expression of eGFP-iPAK4. To infer an onset time for each band, we mapped the onset of eGFP fluorescence relative to the two dye switches. The slope of the plot of inferred eGFP onset time vs nominal PMA addition time was 0.98± 0.10 (mean ± 95% CI, *N* = 34 fibers at 3 h, 32 fibers at 6 h, 40 fibers at 9 h). Extrapolating this fit to *t* = 0 implied that eGFP onset was delayed by 53 minutes relative to the timing of PMA addition (Fig. 4h). We interpret this delay as the time for PMA to activate the cFos promoter plus delays of transcription, translation, and protein folding of eGFP-iPAK4. The standard deviations in the inferred eGFP onset times were, 34 min (3 h), 50 min (6 h) and 34 min (9 h), implying an average absolute timing accuracy of 39 minutes over a measurement with 12 h between fiducial time stamps.

## Discussion

Slowly growing protein assemblies are a promising substrate for massively parallel cellular recordings. The recording strategy relies on three key elements: a protein scaffold, a means to impart fiducial timestamps, and a fluorescent reporter of cellular activity which is irreversibly incorporated into the scaffold during growth. There are opportunities to broaden and improve upon our results along each of these dimensions.

For applications *in vivo*, the most critical need is to modify the protein scaffold so that it does not deform the cells. One approach is to engineer thinner fibers, either by modifying the iPAK4 scaffold to enhance the ratio of axial to radial growth, or by using a different fiber-forming protein which forms thinner fibers.^16^ A further enhancement would be to have pharmacological regulation of the soluble iPAK4 expression level, either by regulation of transcription^29^ or of protein stability^30^, to permit tuning of the temporal dynamic range. Ideally, one would also like a means to trigger fiber nucleation at a defined time, and to have the steady-state monomer concentration only slightly lower than the nucleation concentration so that the period of initial rapid growth is minimized. Finally, one may wish to express both the iPAK4 and the HT-iPAK4 from a single vector, either by making use of endogenous RNA splicing machinery or by applying a partially effective self-cleaving peptide.

At present, the spacing of the fiducial timestamps is constrained to ∼2 h by the ∼4.5 h half-life of soluble HT-iPAK4. If one could shorten this half-life, one could place time stamps closer together, and thereby achieve greater absolute timing accuracy. Examination of the trajectories of soluble and fiber-bound fluorescence during fiber nucleation (Fig. S1b) implies that most soluble iPAK4 is lost via incorporation into the growing fiber. One might shorten the half-life of the soluble HT-iPAK4 by attaching ubiquitination domains or other tags to facilitate proteolytic turnover of the soluble subunits.^31^ Alternatively, one might consider a variety of optogenetic caging strategies to reversibly control the availability of time-stamp monomers.^32^ Optogenetic modulation of protein availability can occur over seconds, suggesting the possibility to introduce extremely precise fiducial time stamps.

Finally, the modular design of iPAK4 fibers could accommodate diverse recording modalities. A simple generalization of the present results would be to record the simultaneous dynamics of multiple promoters by using each to drive expression of iPAK4 fused to a spectrally distinct fluorescent protein. There also exist diverse fluorescent reporters of covalent enzymatic modifications, e.g. of kinase, phosphatase, or protease activity.^33,34^ Such reporters are likely protected from enzymatic modification within the fiber, and thereby might be a means to record the state of the cell at the moment of incorporation into the fiber.

## Supporting information

Supplementary Movie 1: iPAK4 fiber growth

Supplementary Movie 2: Tracking iPAK4 fiber growth

## Acknowledgments

We thank D. Kim, Y. Baskaran and E. Manser for helpful discussions. We thank B. Cui for the PAK4 plasmid and C. Hellriegel at the Harvard Center for BioImaging for assistance with microscopy. We thank S. Begum, A. Preecha, and J. Koob for technical assistance. This work was supported by a Vannevar Bush Faculty Fellowship (AEC), the Howard Hughes Medical Institute (AEC) and the Harvard Brain Initiative (DL).

## Author contributions

DL and AEC conceived the project and designed the experiments. DL and XTL cloned the plasmids. PP and DL performed the patch clamp measurements. XTL performed the time-lapse imaging in HEK293T cells. DL performed all other characterizations in cultured cells and acquired the imaging data. JBG, NF, LDL synthesized and supplied the JF dyes. HS and DB assisted in protein design and optimization. DL, AEC, PP, and BT analyzed the data. DL and AEC wrote the paper. All authors participated in the revision of the manuscript.

## Materials and methods

### Cloning and molecular biology

All iPAK4 constructs were cloned into a second-generation lentiviral backbone (Addgene: 136636) with either a CMV or a cFos promoter (Addgene: 47907) using standard Gibson Assembly.^**35**^ Briefly, the vector was linearized by double digestion (BamHl and EcoRl for CMV driven constructs, Pacl and EcoRl for cFos driven constructs) and purified by the GeneJET gel extraction kit (ThermoFisher). Gene fragments and cFos promoter were generated by PCR amplification and then combined with the linearized backbones by Gibson ligation. EGFP-iPAK4 was a gift from B. Cui (Stanford), and HaloTag was cloned from pCAG-Voltron (Addgene: 119033). The eGFP and iPAK4 were connected with a SGGS linker, while the HaloTag and iPAK4 were connected with a SGS linker. All plasmids were verified by full sequencing around the cloned regions.

All plasmids are available on Addgene:

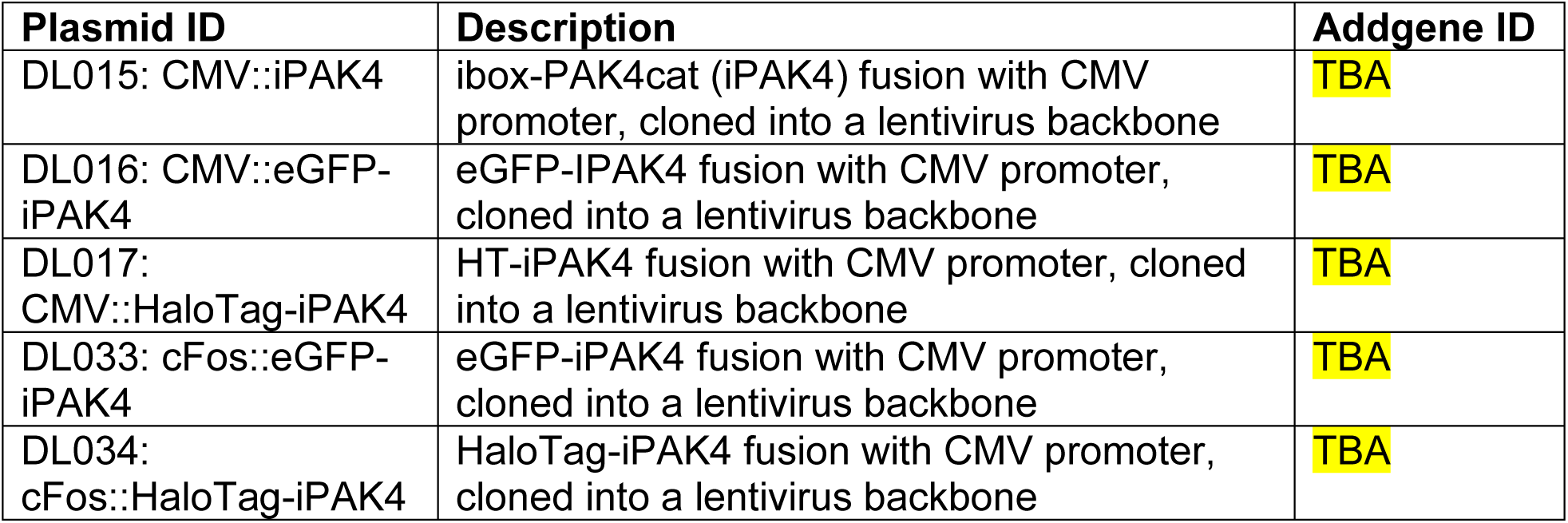

| Plasmid ID | Description | Addgene ID |
| --- | --- | --- |
| DL015: CMV::iPAK4 | ibox-PAK4cat (iPAK4) fusion with CMV promoter, cloned into a lentivirus backbone | TBA |
| DL016: CMV::eGFP-iPAK4 | eGFP-iPAK4 fusion with CMV promoter, cloned into a lentivirus backbone | TBA |
| DL017: CMV::HaloTag-iPAK4 | HT-iPAK4 fusion with CMV promoter, cloned into a lentivirus backbone | TBA |
| DL033: cFos::eGFP-iPAK4 | eGFP-iPAK4 fusion with CMV promoter, cloned into a lentivirus backbone | TBA |
| DL034: cFos::HaloTag-iPAK4 | HaloTag-iPAK4 fusion with CMV promoter, cloned into a lentivirus backbone | TBA |

### Synthesis of JFX_608_-HaloTag

#### General synthetic methods

Commercial reagents were obtained from reputable suppliers and used as received. All solvents were purchased in septum-sealed bottles stored under an inert atmosphere. All reactions were sealed with septa through which a nitrogen atmosphere was introduced unless otherwise noted. Reactions were conducted in round-bottomed flasks or septum-capped crimp-top vials containing Teflon-coated magnetic stir bars. Heating of reactions was accomplished with a silicon oil bath or an aluminum reaction block on top of a stirring hotplate equipped with an electronic contact thermometer to maintain the indicated temperatures. Reactions were monitored by thin layer chromatography (TLC) on precoated TLC glass plates (silica gel 60 F_254_, 250 μm thickness) or by LC/MS (Phenomenex Kinetex 2.1 mm × 30 mm 2.6 μm C18 column; 5 μL injection; 5–98% MeCN/H_2_O, linear gradient, with constant 0.1% v/v HCO_2_H additive; 6 min run; 0.5 mL/min flow; ESI; positive ion mode). TLC chromatograms were visualized by UV illumination or developed with *p*-anisaldehyde, ceric ammonium molybdate, or KMnO_4_ stain. Reaction products were purified by flash chromatography on an automated purification system using pre-packed silica gel columns or by preparative HPLC (Phenomenex Gemini–NX 30 × 150 mm 5 μm C18 column). Analytical HPLC analysis was performed with an Agilent Eclipse XDB 4.6 × 150 mm 5 μm C18 column under the indicated conditions. High-resolution mass spectrometry was performed by the High Resolution Mass Spectrometry Facility at the University of Iowa. NMR spectra were recorded on a 400 MHz spectrometer. ^1^H and ^13^C chemical shifts were referenced to TMS or residual solvent peaks. Data for ^1^H NMR spectra are reported as follows: chemical shift (δ ppm), multiplicity (s = singlet, d = doublet, t = triplet, q = quartet, dd = doublet of doublets, m = multiplet), coupling constant (Hz), integration. Data for ^13^C NMR spectra are reported by chemical shift (δ ppm) with hydrogen multiplicity (C, CH, CH_2_, CH_3_) information obtained from DEPT spectra.

#### 6-tert-Butoxycarbonyl-JFX_608_ (2)

A vial was charged with 6-*tert*-butoxycarbonyl-carbofluorescein ditriflate^20^ (**1**; 250 mg, 0.346 mmol), Pd_2_dba_3_ (31.7 mg, 34.6 μmol, 0.1 eq), XPhos (49.5 mg, 0.104 mmol, 0.3 eq), and Cs_2_CO_3_ (316 mg, 0.969 mmol, 2.8 eq). The vial was sealed and evacuated/backfilled with nitrogen (3×). Dioxane (2 mL) was added, and the reaction was flushed again with nitrogen (3×). Following the addition of azetidine-2,2,3,3,4,4-*d*_6_ (52.4 mg, 0.830 mmol, 2.4 eq), the reaction was stirred at 100 °C for 4 h. It was subsequently cooled to room temperature, filtered through Celite with CH_2_Cl_2_, and concentrated to dryness. Purification by silica gel chromatography (20–100% EtOAc/hexanes, linear gradient) afforded 6-*tert*-butoxycarbonyl-JFX_608_ (**2**) as a blue-green solid (181 mg, 95%). ^1^H NMR (CDCl_3_, 400 MHz) δ 8.14 (dd, *J* = 8.0, 1.3 Hz, 1H), 8.00 (dd, *J* = 8.0, 0.8 Hz, 1H), 7.62 (dd, *J* = 1.2, 0.7 Hz, 1H), 6.58 (d, *J* = 2.4 Hz, 2H), 6.54 (d, *J* = 8.6 Hz, 2H), 6.21 (dd, *J* = 8.6, 2.3 Hz, 2H), 1.83 (s, 3H), 1.73 (s, 3H), 1.53 (s, 9H); ^13^C NMR (CDCl_3_, 101 MHz) δ 170.2 (C), 164.6 (C), 155.6 (C), 152.5 (C), 146.8 (C), 137.8 (C), 130.3 (C), 130.1 (CH), 128.9 (CH), 125.1 (CH), 124.8 (CH), 119.8 (C), 110.5 (CH), 108.0 (CH), 88.8 (C), 82.3 (C), 38.5 (C), 35.5 (CH_3_), 32.8 (CH_3_), 28.2 (CH_3_); Analytical HPLC: t_R_ = 13.3 min, >99% purity (10–95% MeCN/H_2_O, linear gradient, with constant 0.1% v/v TFA additive; 20 min run; 1 mL/min flow; ESI; positive ion mode; detection at 600 nm); HRMS (ESI) calcd for C_34_H_25_D_12_N_2_O_4_ [M+H]^+^ 549.3501, found 549.3503.

#### 6-Carboxy-JFX _608_ (3)

6-*tert*-Butoxycarbonyl-JFX_608_ (**2**; 310 mg, 0.565 mmol) was taken up in CH_2_Cl_2_ (10 mL), and trifluoroacetic acid (2 mL) was added. The reaction was stirred at room temperature for 6 h. Toluene (10 mL) was added; the reaction mixture was concentrated to dryness and then azeotroped with MeOH three times to provide 6-carboxy-JFX_608_ (**3**) as a dark blue solid (323 mg, 94%, TFA salt). Analytical HPLC and NMR indicated that the material was >95% pure and did not require further purification prior to amide coupling. ^1^H NMR (CD_3_OD, 400 MHz) δ 8.36 – 8.27 (m, 2H), 7.86 – 7.78 (m, 1H), 6.91 (d, *J* = 9.1 Hz, 2H), 6.81 (d, *J* = 2.3 Hz, 2H), 6.38 (dd, *J* = 9.1, 2.3 Hz, 2H), 1.82 (s, 3H), 1.70 (s, 3H); ^13^C NMR (CD_3_OD, 101 MHz) δ 168.0 (C), 167.5 (C), 158.0 (C), 157.0 (C), 139.5 (C), 137.6 (CH), 136.2 (C), 135.5 (C), 132.4 (CH), 132.3 (CH), 131.4 (CH), 121.8 (C), 111.9 (CH), 109.7 (CH), 42.8 (C), 35.6 (CH_3_), 32.0 (CH_3_); Analytical HPLC: t_R_ = 10.5 min, >99% purity (10–95% MeCN/H_2_O, linear gradient, with constant 0.1% v/v TFA additive; 20 min run; 1 mL/min flow; ESI; positive ion mode; detection at 600 nm); HRMS (ESI) calcd for C_30_H_17_D_12_N_2_O_4_ [M+H]^+^ 493.2875, found 493.2873.

#### JFX_608_-NHS (4)

6-Carboxy-JFX_608_ (**3**, TFA salt; 125 mg, 0.206 mmol) was combined with DSC (116 mg, 0.453 mmol, 2.2 eq) in DMF (5 mL). After adding Et_3_N (172 μL, 1.24 mmol, 6 eq) and DMAP (2.5 mg, 20.6 μmol, 0.1 eq), the reaction was stirred at room temperature for 30 min. It was subsequently diluted with 10% w/v citric acid and extracted with EtOAc (2×). The combined organic extracts were washed with water and brine, dried over anhydrous MgSO_4_, filtered, and concentrated *in vacuo*. Flash chromatography (25–100% EtOAc/CH_2_Cl_2_, linear gradient) yielded 116 mg (95%) of JFX_608_-NHS (**4**) as a dark blue-green solid. ^1^H NMR (CDCl_3_, 400 MHz) δ 8.30 (dd, *J* = 8.0, 1.4 Hz, 1H), 8.11 (dd, *J* = 8.0, 0.8 Hz, 1H), 7.78 (dd, *J* = 1.4, 0.7 Hz, 1H), 6.57 (d, *J* = 2.4 Hz, 2H), 6.50 (d, *J* = 8.6 Hz, 2H), 6.23 (dd, *J* = 8.5, 2.4 Hz, 2H), 2.87 (s, 4H), 1.83 (s, 3H), 1.71 (s, 3H); Analytical HPLC: t_R_ = 11.1 min, >99% purity (10–95% MeCN/H_2_O, linear gradient, with constant 0.1% v/v TFA additive; 20 min run; 1 mL/min flow; ESI; positive ion mode; detection at 600 nm); HRMS (ESI) calcd for C_34_H_20_D_12_N_3_O_6_ [M+H]^+^ 590.3039, found 590.3043.

#### JFX_608_–HaloTag ligand (6)

JFX_608_-NHS (**4**; 50 mg, 84.8 μmol) and 2-(2-((6-chlorohexyl)oxy)ethoxy)ethanamine (**5**, “HaloTag(O2)amine,” TFA salt; 43.0 mg, 0.127 mmol, 1.5 eq) were combined in DMF (3 mL), and DIEA (44.3 μL, 0.254 mmol, 3 eq) was added. After stirring the reaction at room temperature for 1 h, it was diluted with saturated NaHCO_3_ and extracted with EtOAc (2×). The combined organic extracts were washed with water and brine, dried over anhydrous MgSO_4_, filtered, and concentrated *in vacuo*. Purification of the crude product by silica gel chromatography (50–100% EtOAc/toluene, linear gradient) provided JFX_608_–HaloTag ligand (**6**) as a blue foam (47 mg, 79%). ^1^H NMR (CDCl_3_, 400 MHz) δ 8.02 (dd, *J* = 8.0, 0.7 Hz, 1H), 7.94 (dd, *J* = 7.9, 1.4 Hz, 1H), 7.44 – 7.39 (m, 1H), 6.73 (t, *J* = 4.8 Hz, 1H), 6.57 (d, *J* = 2.4 Hz, 2H), 6.52 (d, *J* = 8.6 Hz, 2H), 6.20 (dd, *J* = 8.6, 2.4 Hz, 2H), 3.64 – 3.56 (m, 6H), 3.54 – 3.48 (m, 4H), 3.38 (t, *J* = 6.6 Hz, 2H), 1.83 (s, 3H), 1.78 – 1.73 (m, 2H), 1.72 (s, 3H), 1.55 – 1.48 (m, 2H), 1.45 – 1.38 (m, 2H), 1.34 – 1.27 (m, 2H); Analytical HPLC: t_R_ = 12.6 min, >99% purity (10–95% MeCN/H_2_O, linear gradient, with constant 0.1% v/v TFA additive; 20 min run; 1 mL/min flow; ESI; positive ion mode; detection at 600 nm); HRMS (ESI) calcd for C_40_H_37_D_12_ClN_3_O_5_ [M+H]^+^ 698.4108, found 698.4118.

### Fiber expression in HEK cells

HEK293T cells were grown and split following standard protocols as described previously.^36^ HEK293T cells at low-passage-number (< 10 passages) were plated at a confluence of 30% onto 10-cm dishes coated with gelatin (Stemcell Technologies; 07903) or 14 mm glass bottom dishes (CellVis, D35-14-1.5-N) coated with 40 μg/ml poly-L-lysine-coated (P8920, Sigma-Aldrich). Dulbecco’s modified Eagle’s medium (DMEM) supplemented with 10% FBS and penicillin/streptomycin was used as the culture medium. Cells were grown at 37 °C and 5% CO_2_.

When cells reached 50-70% confluence, genes were delivered by either lentivirus or TransIT-293 (Mirus; MIR2700) transfection kit. In lentiviral transduction, the high-titer lentiviral vectors were first pre-mixed at the designated ratio and diluted to 10% of the initial concentration by DMEM medium. The cells’ culture medium was then replaced by the fresh lentivirus-containing DMEM medium, and the cells were further incubated at 37 °C and 5% CO_2_ for ∼24 h. In the TransIT-293 transfection, 1 μg plasmids at the designated ratio were first diluted by 100 μl Opti-MEM medium, followed by the addition of 3 μl TransIT-293 reagent. The cocktail was incubated at room temperature for 15 min and diluted 10-fold into DMEM on the culture dishes. The cells were then further incubated at 37 °C and 5% CO_2_ for ∼24 h.

### Fiber labeling

The in-cellulo fibers were labeled with HT-ligand Janelia Fluor (JF) dyes. In this work the cell-permeable dyes were: JF_503_, JF_525_, JF_552_, JFX_608_, and JF_669_. The JF dyes were first diluted into 1 mM stock solution as described previously,_19_ which was aliquoted and stored at −20 °C. The 1 mM solution was further diluted to 1 μM in 37 °C DMEM medium for HEK cells or BPNM/SM1 medium for neurons. The dyed medium was used to replace the original medium in the culture dishes at timed staining. In the dye switching processes, the medium containing the original dye was fully removed, followed by thorough wash of the culture dishes 5 times with 37 °C culture medium. Then, the medium with 1 μM new dye was added to the cells, and the cells were returned to the incubator at 37 °C and 5% CO_2_.

### Measurements of HT dye labeling kinetics in HEK293T cells

HEK293T cells were transfected with an inducible nuclear-localized HT protein (HT-NLS, Addgene #82518). Doxycycline (DOX, 2 μg/ml) was added 12 h after transfection to induce the expression of HT-NLS. 24 h after transfection, cells were incubated with 0.1 μm of the indicated dye and nuclear fluorescence was monitored via wide-field epifluorescence microscopy as a function of time.

### Primary neuron culture

All procedures involving animals were in accordance with the US National Institutes of Health Guide for the care and use of laboratory animals and were approved by the Institutional Animal Care and Use Committee at Harvard University.

Before the plating of primary hippocampal neurons, 14 mm glass-bottom dishes were first incubated with 40 μg/ml poly-D-lysine (PDL) in PBS at room temperature for 1 h and subsequentially with 20 μg/ml laminin (Fisher Scientific; 23-017-015) at 4 °C overnight, followed by thorough wash with PBS. Hippocampi (BrainBits; SKU: SDEHP) from embryonic day 18 (E18) rats were dissected and resuspended in Brainphys_TM_ medium (BPNM, Stemcell Technologies; 05790) supplemented with 2% SM1 (Stemcell Technologies; 05792), 5 mM L-Glutamine (Stemcell Technologies; 07100), and 35 μg/ml L-Glutamic Acid (Sigma Aldrich; 49449), to a final concentration of 3.0 ×10_6_ cells/ml. The neurons were then plated at a density of 30,000 cells/cm_2_ on the pretreated glass-bottom dishes, with subsequent addition of 2 ml BPNM with 2% SM1 (BPNM/SM1). Neuronal health was monitored daily from DIV1 to DIV 7. Every 3-4 days, 1 ml of the medium in each dish was replaced with 37 °C fresh BPNM/SM1 medium.

### Patch clamp electrophysiology

Whole-cell recordings were performed in extracellular buffer containing (in mM): 125 NaCl, 2.5 KCl, 15 HEPES, 25 D-glucose, 1 MgCl_2_, and 2 CaCl_2_ (pH = 7.2-7.3 with NaOH). Fiber-forming and non-forming (control) neurons were visualized with a home-built inverted epifluorescence microscope. Experiments were made at 23 °C under ambient atmosphere. The whole-cell internal solution comprised (in mM): 8 NaCl, 130 KMeSO_3_, 10 HEPES, 5 KCl, 0.5 EGTA, 4 Mg-ATP, and 0.3 Na_3_-GTP. The pH was adjusted to 7.2-7.3 with KOH and osmolarity was set to 290-295 mOsm/L. Borosilicate glass pipettes were used with a resistance of 3-5 MΩ (1.5 mm OD). Signals were acquired and filtered at 4 kHz with the internal Bessel filter using a Multiclamp 700B (Molecular Devices) and digitized with PCIe-6323 (National Instruments) at 10 kHz. Following the whole-cell configuration, membrane capacitance (C_m_), and membrane resistance (R_m_) were estimated under voltage-clamp mode. Measurements of resting membrane potential (V_rest_), rheobase, and spike rates were made under current-clamp mode. Rheobase was defined as the minimum current step (in 500 ms duration) required for any spike onset. Whole-cell recordings were monitored and analyzed using a custom code written in LabView and Matlab.

### Lentivirus production

#### Lentivirus production in HEK293T cells

Plasmids of CMV::iPAK4, CMV::HaloTag-iPAK4, and cFos::eGFP-iPAK4 were used to produce lentivirus according to published methods._37_ Briefly, low passage-number HEK293T cells (ATCC; CRL-11268) were plated onto gelatin-coated (Stemcell Technologies; 07903) 10-cm dishes. When HEK cells reached 80% confluence, the medium was exchanged to a serum-free DMEM. After 0.5–1 h, cells were transfected using polyethylenimine (PEI; Sigma; 408727). 7 μg of the vector plasmid, 4 μg of the second-generation packaging plasmid psPAX2 (Addgene; 12260), and 2 μg of viral entry protein VSV-G plasmid pMD2.G (Addgene; 12259) were mixed into 600 μl of serum-free DMEM, and 20 μl of 1 mg/ml PEI was then added. The mixture was incubated at room temperature for 15 min and added dropwise to the plate. After 3-4 h, the medium was exchanged back to 10 ml of DMEM10. The supernatant was harvested at 36 h post-transfection, and another 10 ml of DMEM10 was added to the cells and incubated for another 24 h. At 60 h post-transfection, the supernatant was harvested again and combined with the first batch of supernatant, centrifuged for 5 min at 500 g, and filtered through a 0.45-μm filter (EMD Millipore; SE1M003M00).

#### Lentivirus concentration

1 part of Lenti-X™ concentrator (TaKaRa; 631232) was first mixed with 3 parts of supernatant and incubated at 4 °C overnight for lentivirus precipitation. The mixture was then centrifuged at 1,500 g for 45 min at 4°C. The supernatant was gently removed, and the off-white pellet was resuspended in 200 μl neurobasal-based medium. The concentrated virus was titrated in neurons, aliquoted, and stored at −80 °C for neuronal transduction.

#### Fiber expression in neurons

Genes were delivered to neurons by lentiviral transduction at DIV 7-10. The lentiviral vectors of CMV::iPAK4, CMV::HaloTag-iPAK4, and cFos::eGFP-iPAK4 were first mixed at the designated ratio, which was further diluted to 10% of the original concentration by fresh BPNM/SM1 medium. The dilution was then used to replace the original medium in neuronal culture. The neurons were incubated in the lentivirus-containing medium at 37 °C and 5% CO_2_ for 12 h, followed by medium replacement with lentivirus-free medium.

#### Chemical activation of neuronal activity

The cFos promoter was activated by phorbol 12-myristate 13-acetate (PMA; Sigma Aldrich P8139). Briefly, the PMA was first diluted with DMSO to form a 1 mM stock solution. At the designated time, 2 μl PMA stock was directly added to each 14 mm glass-bottom culture dish, which contained 2 ml BPNM/SM1 medium. The dishes were then stirred gently to mix the PMA and medium. After PMA addition, the dishes were returned to an incubator at 37 °C with 5% CO_2_.

#### Multispectral imaging

Multispectral images were acquired using ZEISS LSM 980 confocal microscope with Airyscan 2. Lambda scan mode was used to image fibers with multi-color labeling. The excitation laser wavelengths were 488 nm (eGFP, JF_503_, JF_525_), 561 nm (JF_552_, JFX_608_), and 639 nm (JF_669_). In each Lambda scan, 32 channels in the range of 414-688 nm were simultaneously acquired to obtain a hyperspectral stack of images. The images were then unmixed with the built-in linear unmixing algorithm in Zen Blue software. Reference images of individual fluorescent labels were taken in the same instrumental configuration to train the linear unmixing algorithm. The spectral unmixing typically produced negligible residual signals.

#### Time-lapse microscopy

HEK293T cells expressing the target constructs were grown on 14 mm glass bottom culture dishes (CellVis, D35-14-1.5-N) and were monitored under a Zeiss Elyra microscope with 488 nm laser and a 10x air objective in an environmental chamber at 37 °C and 5% CO_2_. Images were acquired at 1% laser power, 300 ms exposures, 10 min intervals over 10-23 hours post-transfection.

#### Image processing and data analysis

Images of individual fibers were rotated to align the long axis to the x-axis. Fluorescence profiles were then calculated as the median fluorescence in each spectral channel across the width of the fiber. Fibers which were not in focus or where two or more fibers crossed each other near a dye transition were excluded from analysis. Dye transitions were identified as local maxima in the second derivative of the dye fluorescence as a function of position. To avoid spurious peaks due to noise, the second derivative signal was smoothed with a kernel of typically 10 min, though this smoothing was omitted when calculating the width of the dye transition (Fig. S3a).

For tracking fibers during time-lapse recordings, a region of interest (ROI) was manually defined on a maximum intensity projection of the image stack, to select individual fibers and to encompass the entire fiber in all frames of the movie. A Radon transform was then calculated on the selected ROI for each video frame. The peak of the Radon transform was associated with the fiber. The corresponding line in the real-space movie was used to calculate the fiber intensity profile. Nearby parallel lines on either side of the fiber were averaged and used for background subtraction. The fiber ends were then found by applying a simple threshold to the plot of fluorescence vs. position. The fluorescence of the cytoplasm was determined by summing the intensity from pixels that were on-cell but off-fiber.

