## Supplementary Figures for "Time-tagged ticker tapes for intracellular recordings"

**Supplementary Movie Captions**

**Movie S1. Time-lapse recording of iPAK4 fiber growth.** HEK293T cells co-expressed CMV::iPAK4 (95%) and CMV::eGFP-iPAK4 (5%). Time-lapse video microscopy was acquired over a 13 h interval starting 10 h after transfection, with one frame every 10 min.

**Movie S2. Tracking individual iPAK4 fibers in HEK293T cells.** Left: The time-lapse movie was manually segmented to define regions of interest (ROIs) around individual cells. ROIs were analyzed by Radon transform and the peak of the Radon transform was used to identify the position and orientation of the fiber. Center: Line corresponding to the peak of the Radon transform. Right: A fiber fluorescence profile was calculated from the fluorescence along the maximum intensity line. Mean fluorescence from parallel lines ~1 μm to either side was used for background subtraction. A simple threshold was applied to determine fiber length.

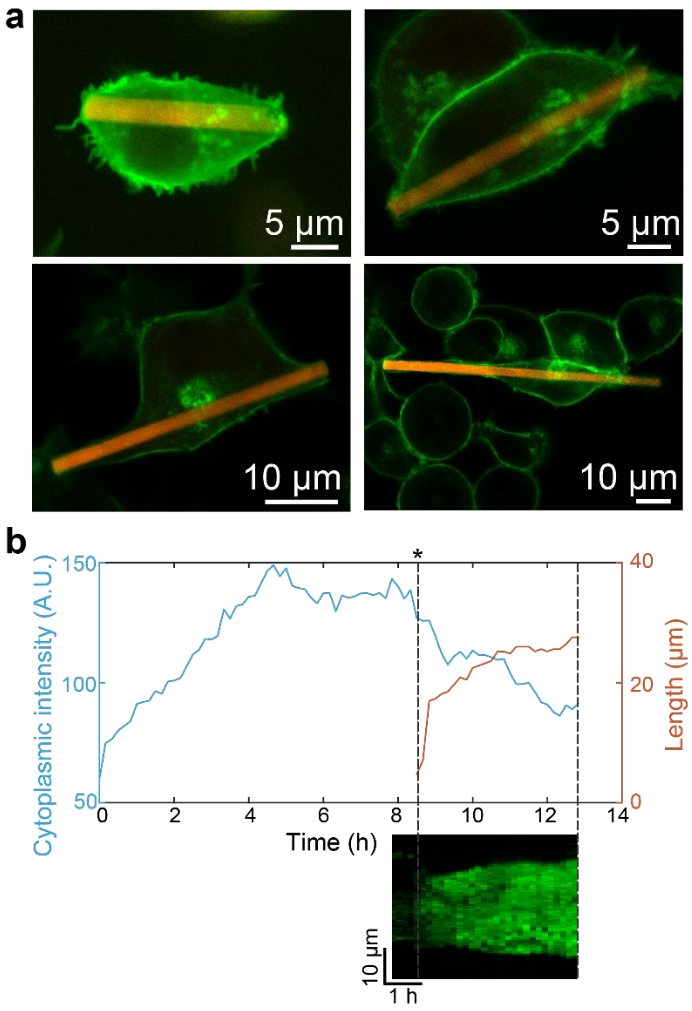


**Figure S1. iPAK4 fiber morphology, nucleation and growth.** Related to Fig. 1. a) Images of HEK293T cells expressing CMV::iPAK4 (95%) and CMV::HT-iPAK4 (5%). The fibers were stained with $JFX_{608}$ and the membrane was labeled by expressing GPI-eGFP. In cells where the fiber length exceeded the cell size, the membrane wrapped around the extended fiber. b) Nucleation and growth of a single eGFP-labeled iPAK4 fiber. HEK293T cells co-expressed CMV::iPAK4 (95%) and CMV::eGFP-iPAK4 (5%). Time-lapse video microscopy was acquired over a 13 h interval starting 10 h after transfection, with one frame every 10 min. Fibers were tracked using a Radon transform algorithm (Methods). The fiber nucleated ~8.5 h after the start of the recording, grew rapidly for the first ~20 min, and then slowed. Fiber nucleation and initial growth were accompanied by a decrease in cytoplasmic eGFP fluorescence. Bottom: Kymograph showing the fiber profile. See also Supplementary Movies 1 and 2.

**a**

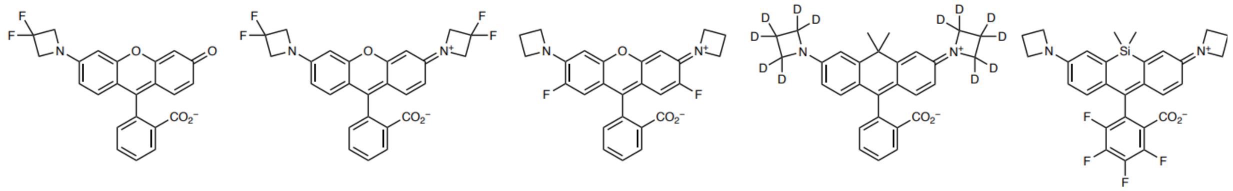

| | $JF_{503}$ | $JF_{525}$ | $JF_{552}$ | $JFX_{608}$ | $JF_{669}$ |
|---|---|---|---|---|---|
| $\lambda_{abs}$ (nm) | 503 | 525 | 552 | 608 | 669 |
| $\varepsilon$ ($M^{-1}cm^{-1}$) | 83,000 | 94,000 | 95,000 | 111,000 | 112,000 |
| $\lambda_{em}$ (nm) | 529 | 549 | 575 | 628 | 682 |
| $\Phi_f$ | 0.87 | 0.91 | 0.83 | 0.74 | 0.37 |

**b**

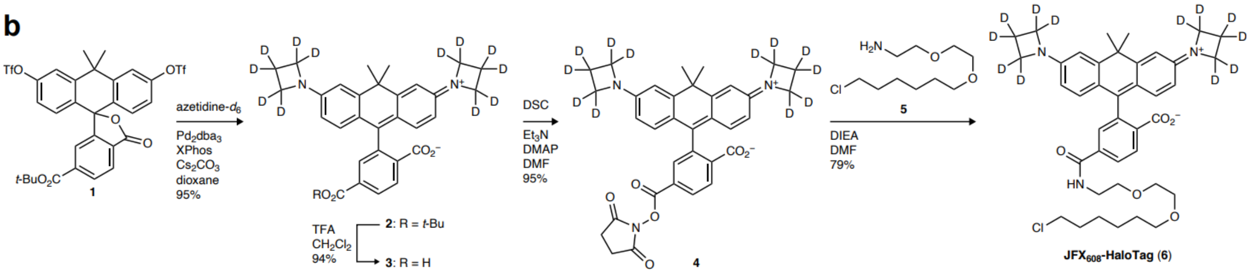


**Figure S2. JaneliaFluor HaloTag ligand dyes used in this study.** a) Chemical structures and spectral properties of the free Janelia Fluor dyes: $JF_{503}$, $JF_{525}$, $JF_{552}$, $JFX_{608}$, and $JF_{669}$. b) Synthesis of $JFX_{608}$-HaloTag.

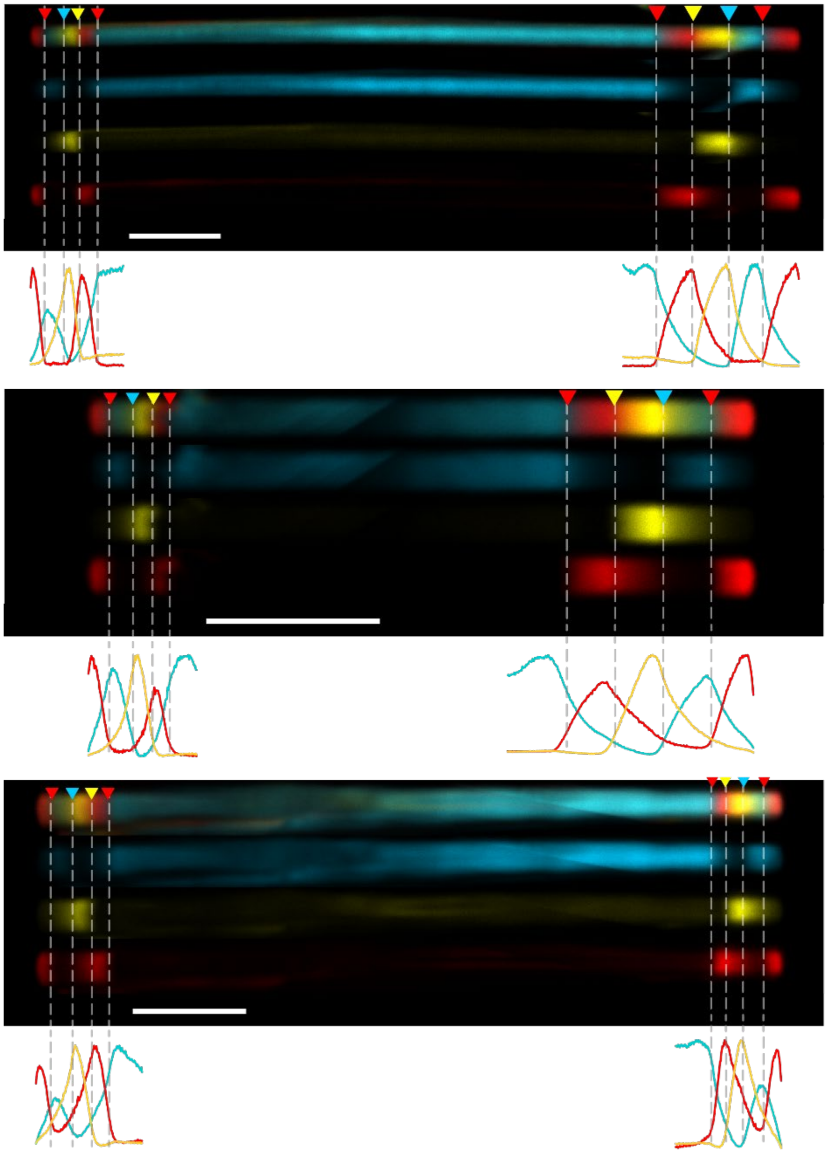

**Figure S3. Color stripes in sequentially labeled HT-iPAK4 fibers.** Related to Fig. 2. Images of HT-iPAK4 fibers labeled with four dye transitions on each end. HEK293T cells were co-transfected with CMV::iPAK4 (95%) and CMV::HT-iPAK4 (5%). After the onset of fiber growth, cells were washed at $\Delta t$ = 2 h intervals in the sequence $JF_{503}$, $JF_{669}$, $JFX_{608}$, $JF_{503}$, $JF_{669}$. Each image panel shows a composite image of a fiber (top), the three-color channels individually (middle), and the line-profiles through each color channel (bottom). Blue: $JF_{503}$, Yellow: $JFX_{608}$, Red: $JF_{669}$. Scale bars 10 $\mu$m.

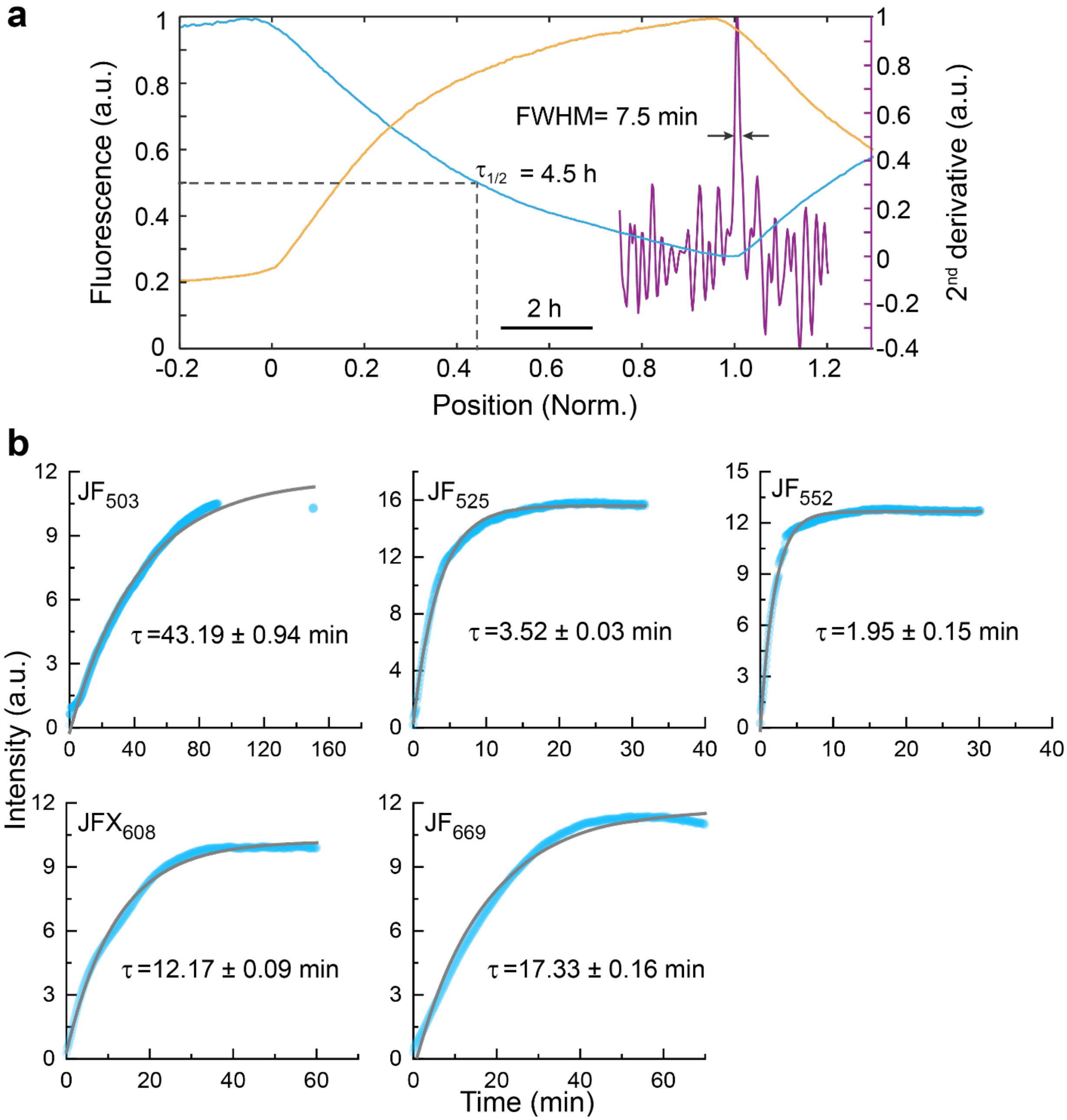


**Figure S4. Kinetics of HT dye labeling and turnover in HEK293T cells.** Related to Fig. 2. a) Mean fluorescence profile of fibers with dye switches at $t$ = 0 and 10 h ($N$ = 200 fibers, length normalized to 0 – 1). The half-life of soluble labeled HT-iPAK4 was 4.5 h, set by the rate of incorporation into the growing fiber. The sharpness of the dye transition was 7.5 min, as calculated by the width of the peak in the second derivative of fluorescence vs position (purple). b) Kinetics of HT dye reaction in HEK293T cells. HEK293T cells expressed a nucleus-targeted HT construct (HT-NLS, Addgene #82518) were incubated with 1 μM of the indicated dye and nuclear fluorescence was monitored as a function of time.

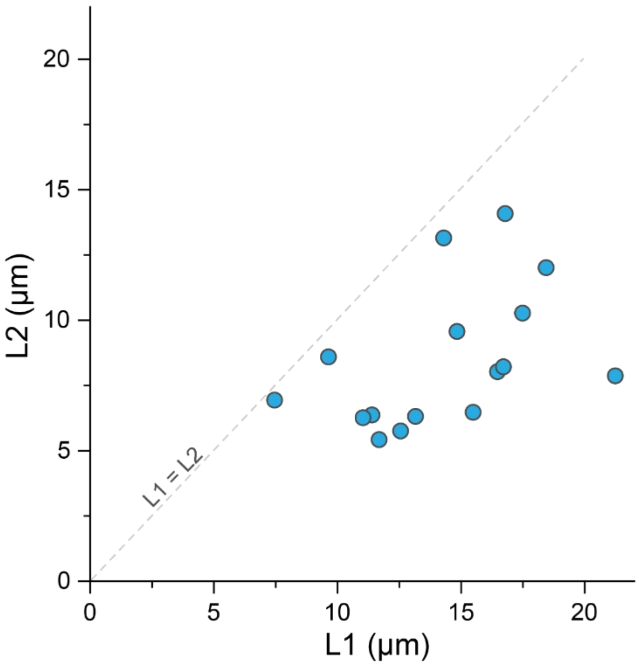


**Figure S5. Comparison of growth rates on the faster and slower-growing fiber ends.** Related to Fig. 2. Dye switches were used to demarcate an 8 h interval on HT-iPAK4 fibers. Both fiber ends were imaged. The longer end was labeled L1, the shorter end L2.

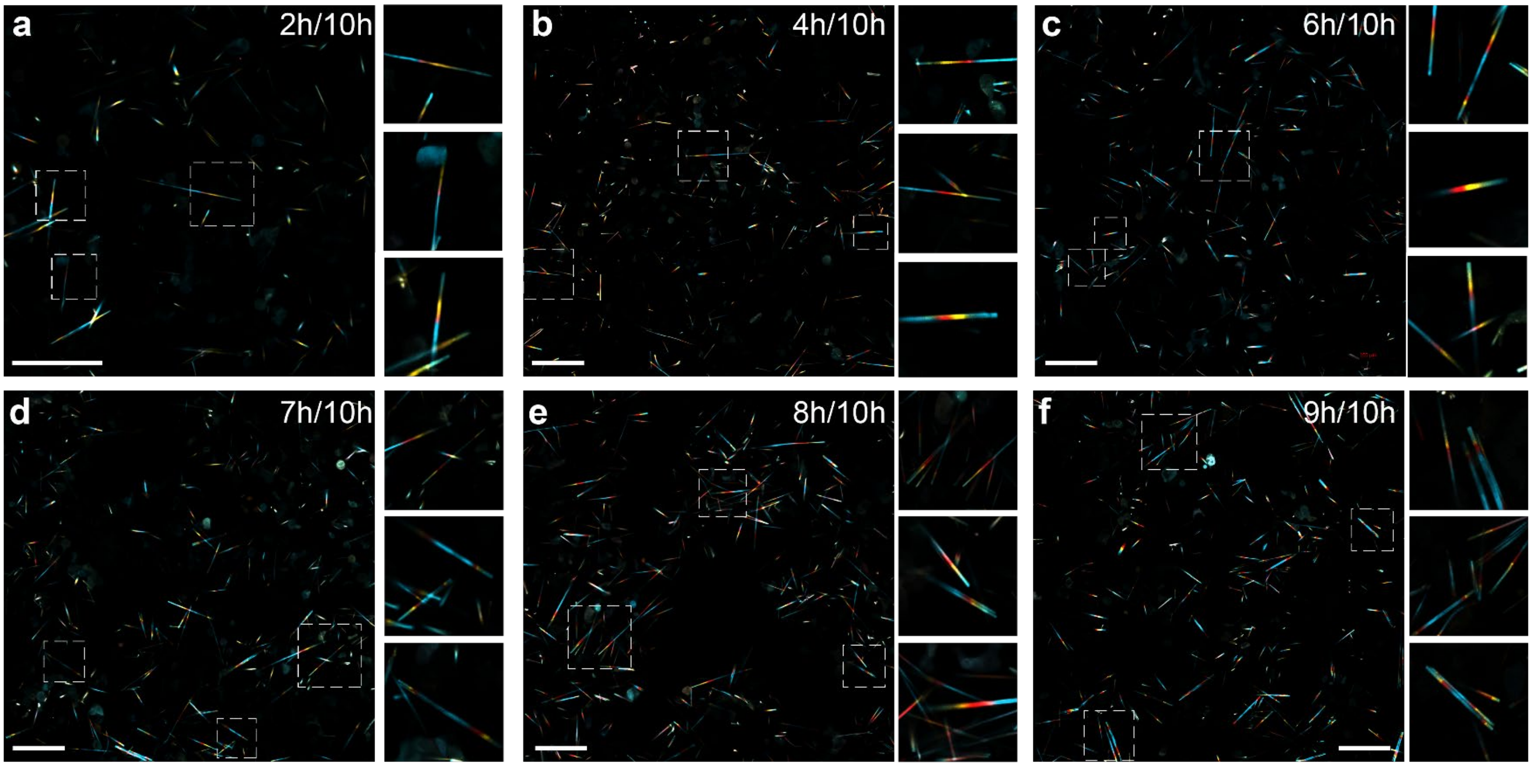


**Figure S6. Low-magnification images of HT-iPAK4 fibers with multiple fiducial timestamps.** Related to Fig. 3. Images show fibers initially labeled with $JF_{525}$, switched to $JF_{669}$ at $t$ = 0, $JFX_{608}$ at $t$ = $X$, and back to $JF_{525}$ at $t$ = 10 h, for $X$ = 2, 4, 6, 7, 8, and 9 h. Insets show the regions in the dashed boxes. Scale bars 100 μm.

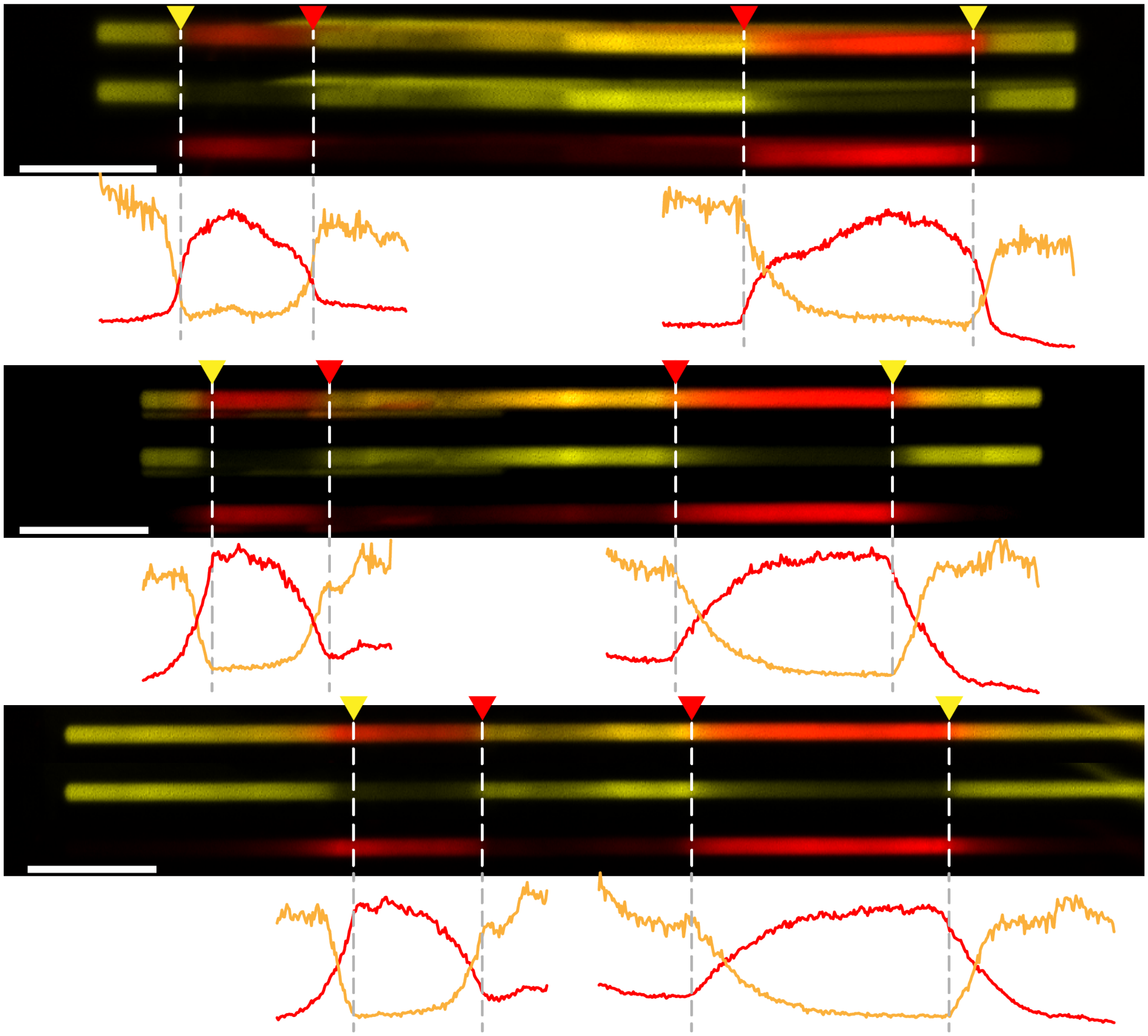

**Figure S7. Fiducial timestamps in neurons expressing HT-iPAK4.** Related to Fig. 4. Neurons were co-infected with lentivirus encoding CMV::iPAK4 (90%) and CMV::HT-iPAK4 (10%). Neurons were initially labeled with $JFX_{608}$ for 24 h, then with $JF_{669}$ for 24 h, and then with $JFX_{608}$ for another 24 h. Images show individual fibers (top), the $JFX_{608}$ (yellow) and JF669 (red) color channels separately (middle), and fluorescence profiles of each color channel (bottom). Scale bars 10 μm.

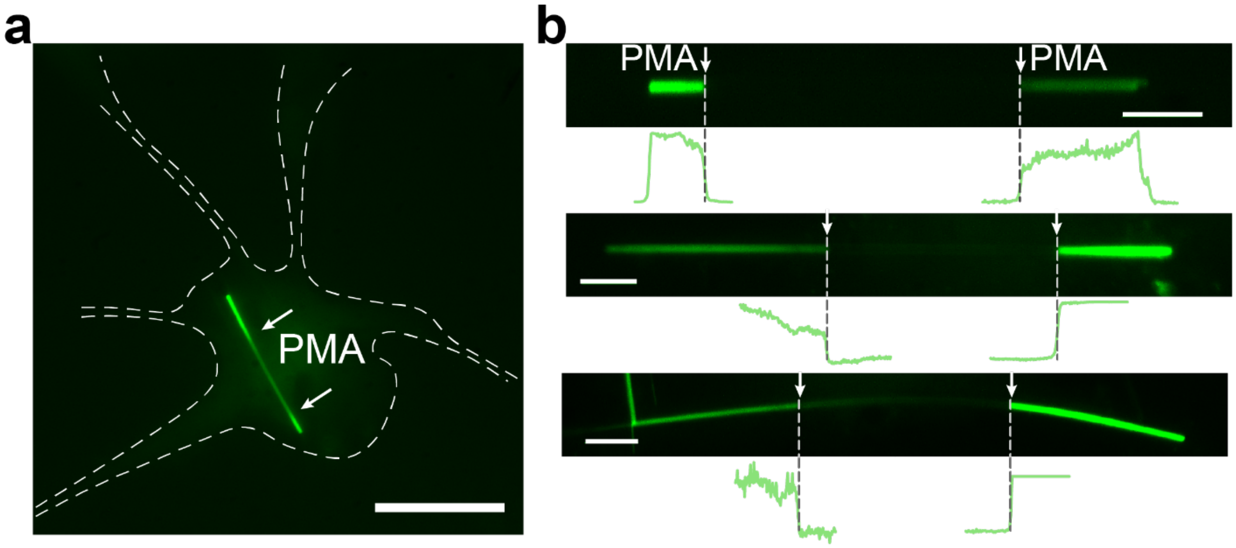


**Figure S8. Monitoring cFos activation in neurons.** Related to Fig. 4. In neurons expressing CMV::iPAK4 (90%) and cFos::eGFP-iPAK4 (10%), addition of PMA activated the *cFos* promoter and led to formation of green stripes in the fibers. a) Example of a fiber inside a neuron. The outline of the neuron is shown with a dotted line. Scale bar 20 μm. b) Examples of individual fibers showing onset of eGFP fluorescence upon addition of PMA. Scale bars 5 μm
