## Supplementary figures and images for "Time-tagged ticker tapes for intracellular recordings"

### Supplementary Movie 1: iPAK4 fiber growth

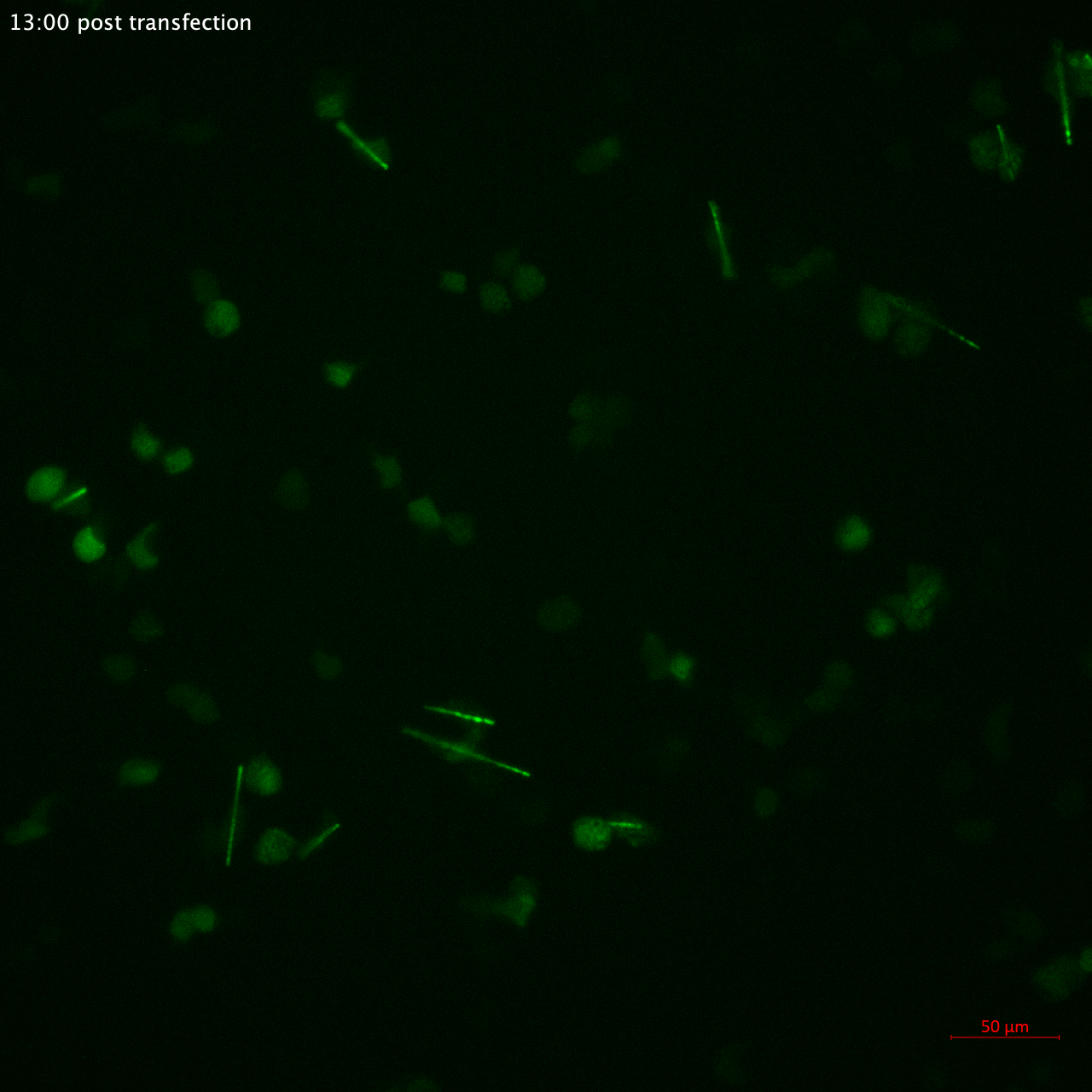
